# BldC delays entry into development to produce a sustained period of vegetative growth in *Streptomyces venezuelae*

**DOI:** 10.1101/194126

**Authors:** Matthew J. Bush, Govind Chandra, Mahmoud M. Al-Bassam, Kim C. Findlay, Mark J. Buttner

## Abstract

Streptomycetes are filamentous bacteria that differentiate by producing spore-bearing reproductive structures called aerial hyphae. The transition from vegetative to reproductive growth is controlled by the *bld* (bald) loci, and mutations in *bld* genes prevent the formation of aerial hyphae, either by blocking entry into development (typically mutations in activators) or by inducing precocious sporulation in the vegetative mycelium (typically mutations in repressors). One of the *bld* genes, *bldC*, encodes a 68-residue DNA-binding protein related to the DNA-binding domain of MerR-family transcription factors. Recent work has shown that BldC binds DNA by a novel mechanism, but there is less insight into its impact on *Streptomyces* development. Here we used ChIP-seq coupled with RNA-seq to define the BldC regulon in the model species *Streptomyces venezuelae*, showing that BldC can function both as a repressor and as an activator of transcription. Using electron microscopy and time-lapse imaging, we show that *bldC* mutants are bald because they initiate development prematurely, bypassing the formation of aerial hyphae. This is consistent with the premature expression of BldC target genes encoding proteins with key roles in development (e.g. *whiD*, *whiI*, *sigF*), chromosome condensation and segregation (e.g. *smeA-sffA*, *hupS*), and sporulation-specific cell division (e.g. *dynAB*), suggesting that BldC-mediated repression is critical to maintain a sustained period of vegetative growth prior to sporulation. We discuss the possible significance of BldC as an evolutionary link between MerR family transcription factors and DNA architectural proteins.

**IMPORTANCE:** Understanding the mechanisms that drive bacterial morphogenesis depends on the dissection of the regulatory networks that underpin the cell biological processes involved. Recently, *Streptomyces venezuelae* has emerged as an attractive new model system for the study of morphological differentiation in *Streptomyces*. This has led to significant progress in identifying the genes controlled by the transcription factors that regulate aerial mycelium formation (Bld regulators) and sporulation (Whi regulators). Taking advantage of *S. venezuelae*, we used ChIP-seq coupled with RNA-seq to identify the genes directly under the control of BldC. Because *S. venezuelae* sporulates in liquid culture, the complete spore-to-spore life cycle can be examined using time-lapse microscopy, and we applied this technique to the *bldC* mutant. These combined approaches reveal BldC to be a member of an emerging class of Bld regulators that function principally to repress key sporulation genes, thereby extending vegetative growth and blocking the onset of morphological differentiation.

## INTRODUCTION

The complex *Streptomyces* life cycle involves two distinct filamentous cell forms: the growing or vegetative hyphae and the reproductive or aerial hyphae, which differentiate into long chains of spores (1-6). Genetic studies identified the regulatory loci that control entry into development, which are called *bld* (bald) genes because null mutations in these loci prevent the formation of aerial hyphae. However, baldness can arise for two different reasons. The larger class of *bld* mutants, which define positive regulators, fail to initiate development, forming colonies of undifferentiated vegetative mycelium. In contrast, a smaller but growing class of *bld* mutants, which define negative regulators, enter development prematurely, inducing sporulation in the vegetative mycelium and bypassing the formation of aerial hyphae. Thus, macroscopically these two classes of mutants look similar, forming smooth colonies that lack the ‘hairy’ appearance of the wild type, but microscopically it is apparent that they arise for diametrically opposed reasons (5, 7-9).

BldC is a small, 68-residue protein with a winged Helix-Turn-Helix (wHTH) motif, related to those found in MerR-family proteins (10). The basic structure of classical MerR proteins is a dimer consisting of two identical subunits, each composed of an N-terminal wHTH DNA-binding domain, a C-terminal effector-recognition domain and an interconnecting linker region that consists of a long α-helix that interacts with the same helix in the other subunit, forming an antiparallel coiled-coil responsible for homodimerization. MerR proteins share significant sequence similarity only within their DNA-binding domains; as different family members bind different effectors, their C-terminal domains are variable and show little, if any, similarity to one another.

MerR transcription factors bind to palindromic DNA sequences as homodimers. However, unlike classical members of the MerR family, BldC has neither an effector domain nor the dimerization helix, and BldC behaves as a monomer in free solution (11). As a consequence, how BldC might bind DNA remained unclear. To address this question, Schumacher et al. (11) carried out biochemical and structural studies to characterize the binding of *S. coelicolor* BldC to the promoters of two known target genes, *whiI* and *smeA*. These studies showed that BldC binds DNA in a completely different way to classical MerR regulators, instead involving asymmetric, cooperative, head-to-tail oligomerization on DNA direct repeats with concomitant pronounced DNA distortion (11). The number of direct repeats present in BldC-binding sites is variable, thus allowing cooperative, head-to-tail binding of additional BldC monomers. Since BldC-like proteins radiate throughout the bacteria, this study identified BldC as the founding member of a new structural family of transcription factors.

Although the work by Schumacher et al. (11) has provided a clear mechanistic understanding of how BldC binds DNA, there has been less insight into its biological role and impact on *Streptomyces* development. In part, this is because previous studies have focussed on the classical model species, *S. coelicolor*, which sporulates only on solid medium. Here we exploit the benefits of the new model species, *Streptomyces venezuelae*, which sporulates in liquid culture (12), to study the biological role of BldC. Using ChIP-seq coupled with RNA-seq, we identify the genes under BldC control and show that BldC can function both as a repressor and as an activator of transcription. We show that *bldC* mutants are bald because they enter development prematurely, bypassing the formation of aerial hyphae. This correlates with the premature expression of BldC target genes with key roles in development, chromosome condensation and segregation, and sporulation-specific cell division, suggesting that BldC-mediated repression is critical to maintain a sustained period of vegetative growth prior to sporulation.

## RESULTS

### Deletion of *bldC* causes premature initiation of development

We constructed an *S. venezuelae bldC* mutant by replacing the *bldC* coding region with an apramycin resistance (*apr*) cassette. The resulting mutant was bald, unable to produce the reproductive aerial hyphae that give mature wild-type *Streptomyces* colonies their characteristic fuzzy appearance (Fig. 1). However, scanning electron microscopy (SEM) of mature colonies of the *bldC* mutant showed that most of the biomass consisted of spores, rather than undifferentiated vegetative hyphae (Fig. 2). Comparison of the growth of the wild type and the *bldC* mutant on plates over time showed that after one day they looked similar (vegetative growth only) but after two days the wild type had produced aerial hyphae while the *bldC* mutant was still restricted to vegetative growth. After 3 days, the aerial hyphae of the wild type had differentiated into spores, and most of the biomass of the *bldC* mutant had also differentiated into spores, bypassing aerial mycelium formation. The *bldC* mutant also seemed to produce higher levels of extracellular matrix than the wild type (Fig. 2). The *bldC* mutant phenotype was fully complemented by introducing a single copy of the *bldC* gene under the control of its native promoter, expressed *in trans* from the ΦBT1 integration site (Figs. 1 and 2).

**FIG 1.**
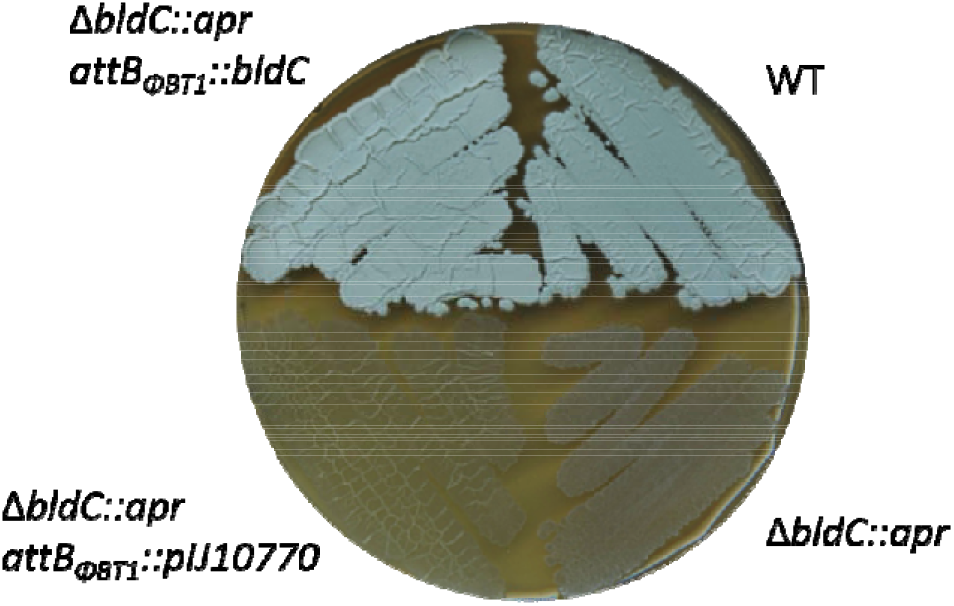
BldC is required for the formation of aerial mycelium. Wild-type *S.venezuelae*, the *bldC* mutant, the *bldC* mutant carrying the empty vector, and the complemented *bldC* mutant, photographed after four days of growth on MYM solid medium.

**FIG 2.**
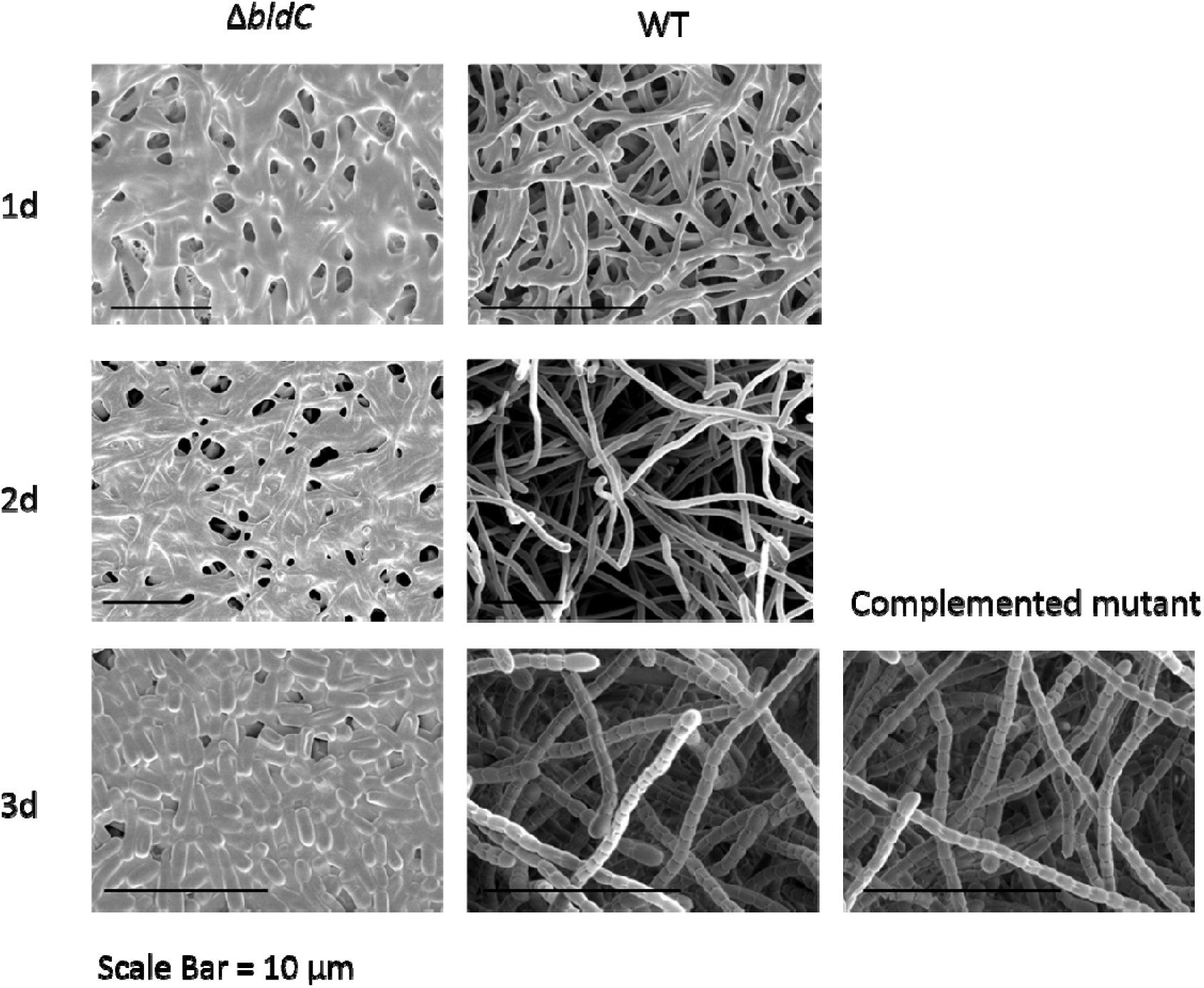
Deletion of *bldC* causes premature initiation of development on solid medium. Scanning electron micrographs showing the phenotypes of the *bldC* mutant and the wild type after one, two and three days of growth on MYM solid medium. The phenotype of the complemented *bldC* mutant is also shown after 3 days of growth on MYM solid medium.

Using an established microfluidic system and methodology (12), we conducted fluorescence time-lapse microscopy to further study the developmental defects associated with deletion of *bldC*. As in previous studies (7, 12-13), we introduced an FtsZ-YPet translational fusion into the wild type, mutant and complemented mutant strains, allowing us to monitor each of the two distinct modes of cell division that occur in *Streptomyces*. In Fig. 3, the scattered single Z-rings mark the position of vegetative cross-walls, which do not constrict or give rise to cell-cell separation, but simply divide the vegetative hyphae into long, box-like compartments (e.g. Figs. 3A + C, panel 2). In contrast, during reproductive growth, long ladders of regularly spaced Z-rings are synchronously deposited along sporogenic hyphae. These Z-rings mark the sites of sporulation septa, which do constrict, ultimately leading to the formation of chains of spores (e.g. Figs. 3A and C, panels 3 and 4). Time-lapse imaging of strains harbouring the FtsZ-YPet fusion showed that the duration of vegetative growth was shorter in the *bldC* mutant compared to the wild type and the complemented mutant (Fig. 3 and Movies S1 A/B, S2 A/B and S3 A/B). Noticeably, following germination, hyphal outgrowth in the *bldC* mutant was associated with an immediate increase in FtsZ-YPet expression, leading to the precocious formation of ladders of Z-rings (Fig. 3B and Movie S2A/B). However, although ladders of Z-rings were observed as early as 4 hours in the *bldC* mutant, mature spores were not observed in the corresponding DIC images until 21 hours, the same time that mature spores were seen in the wild type (Figs. 3A and B). Wild-type patterns of FtsZ expression and sporulation were restored in the complemented mutant (Fig. 3C and Movie S3A/B). From these data, we concluded that the overall role of BldC is to sustain vegetative growth and delay entry into development.

**FIG 3.**
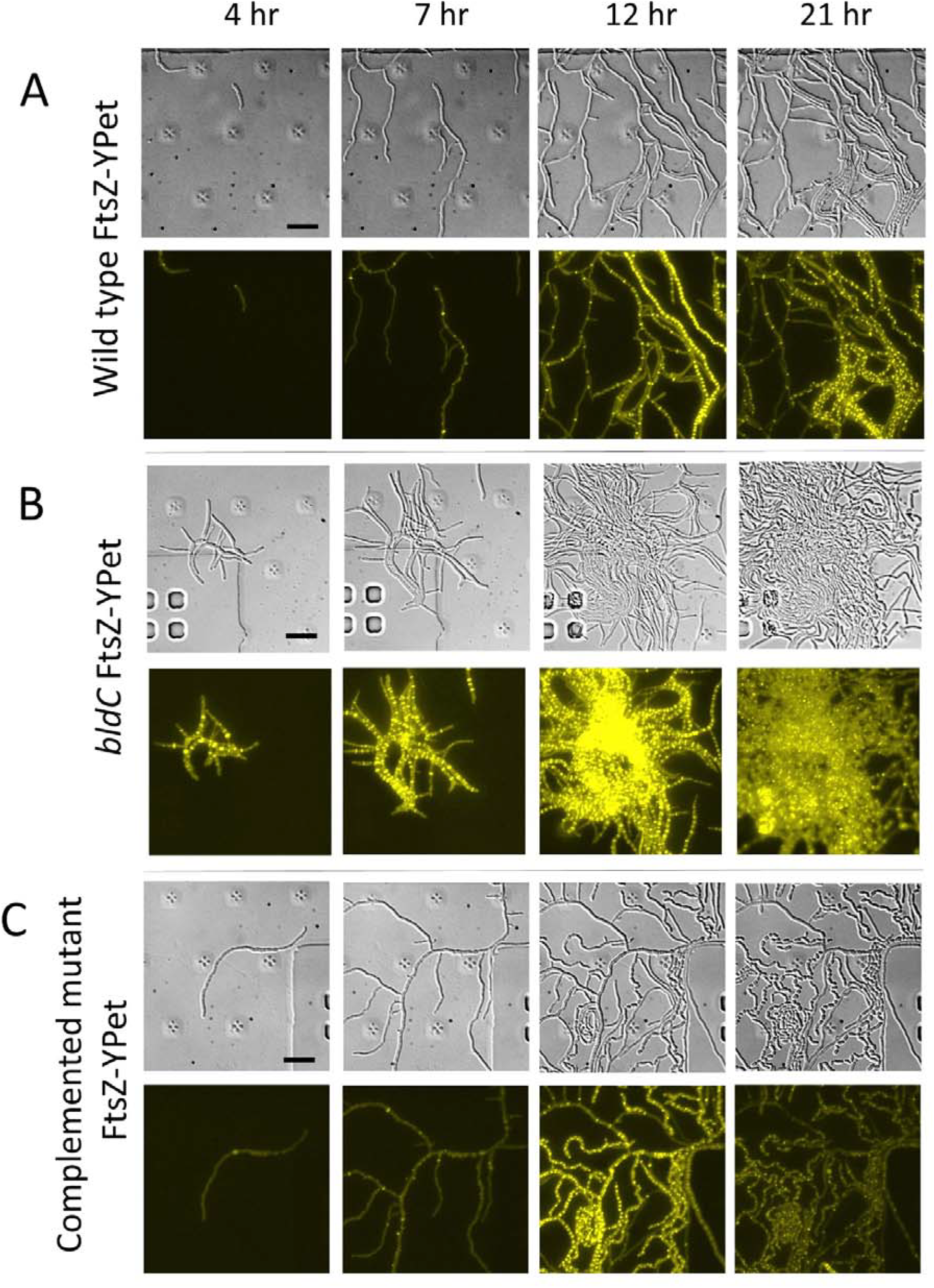
Deletion of *bldC* causes premature initiation of development in liquid medium. Time-lapse images (4, 7, 12 and 21 h post-inoculation) of (A) wild-type *S. venezuelae*, (B) the *bldC* mutant, and (C) the complemented *bldC* mutant, grown in liquid MYM medium in the microfluidic system. All three strains carry the same FtsZ-YPet translational fusion expressed from the native ftsZ promoter, and both the DIC (upper) and fluorescence (lower) images are shown. Scale Bar = 10μm. For the corresponding movies, please see Supporting Information Movies S1A/B, S2A/B and S3 A/B.

### BldC levels are highest early in development

Using an anti-BldC polyclonal antibody, we monitored BldC levels in *S. venezuelae* during sporulation in liquid culture. Western blotting showed that BldC is abundant throughout the life cycle, but that BldC levels are highest early on, during vegetative growth (Fig. 4).

**FIG 4.**
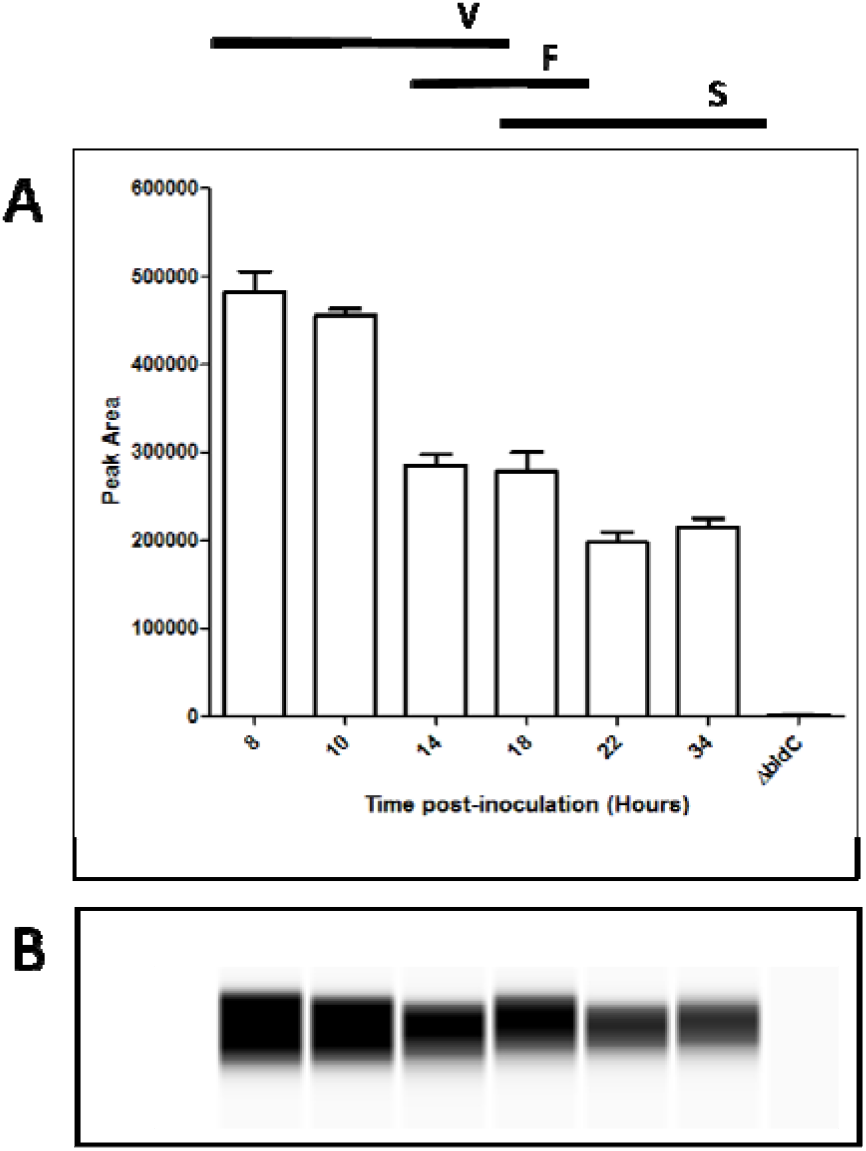
Automated Western blot analysis of BldC levels during submerged sporulation in MYM liquid medium. Equal amounts (1 μg) of total protein were loaded for each sample and BldC was detected with a polyclonal antibody using the quantitative ‘Wes’ capillary electrophoresis and blotting system (ProteinSimple – San Jose, CA). The *S. venezuelae bldC* mutant was used as a negative control. (A) quantitation of BldC levels (area under each peak; arbitrary units). (B) virtual Western blot. All experimental samples were analysed in triplicate and the mean value and its Standard Error are shown for each sample. Each time point is indicated in hours, along with its relation to the developmental stage (V = vegetative growth; F = fragmentation; S = sporulation), as determined by microscopy. Cultures used to analyse BldC levels were identical to those used to prepare RNA prior to qRT-PCR analysis (Fig. 6).

### Defining the BldC regulon in *S. venezuelae*

Previously, ChIP-seq (or ChIP-chip) coupled with transcriptional profiling has enabled us to define the regulons of several key developmental regulators in *S. venezuelae* (14-17). Here, we employed the same approach, using an anti-BldC polyclonal antibody to identify the promoters directly bound by BldC. We performed ChIP-seq at two distinct stages of vegetative growth when BldC was abundant: early vegetative growth (10 hr) and pre-sporulation (14 hr). This work revealed ~360 potential gene targets, the majority of which were bound by BldC at both time points (Table S1A). These targets include many genes encoding key transcriptional regulators of the *Streptomyces* developmental cascade (e.g. *bldM*, *whiB*, *wblA, whiD*, *whiH*, *whiI*, *sigF* and *bldC* itself), others encoding proteins involved in chromosome condensation and segregation during sporulation (e.g. *hupS*, *smeA-sffA*), and those directly involved in cell division during sporulation (e.g. *dynAB, ssgB*) (Fig. 5 and Table 1). Schumacher et al. (11) characterized the interaction of *S. coelicolor* BldC with the promoters of two of its previously known targets, *whiI* and the *smeA-ssfA* operon. *whiI* encodes an orphan response regulator that is essential for the later stages of sporulation, when it forms a functional heterodimer with a second orphan response regulator, BldM, enabling WhiI to bind to DNA and regulate the expression of ~40 sporulation genes (14). The *smeA-sffA* operon encodes a small membrane protein (SmeA) that recruits a DNA translocase (SffA) to sporulation septa (18). Deletion of *smeA-sffA* results in a defect in spore chromosome segregation and has pleiotropic effects on spore maturation (18).

**FIG 5.**
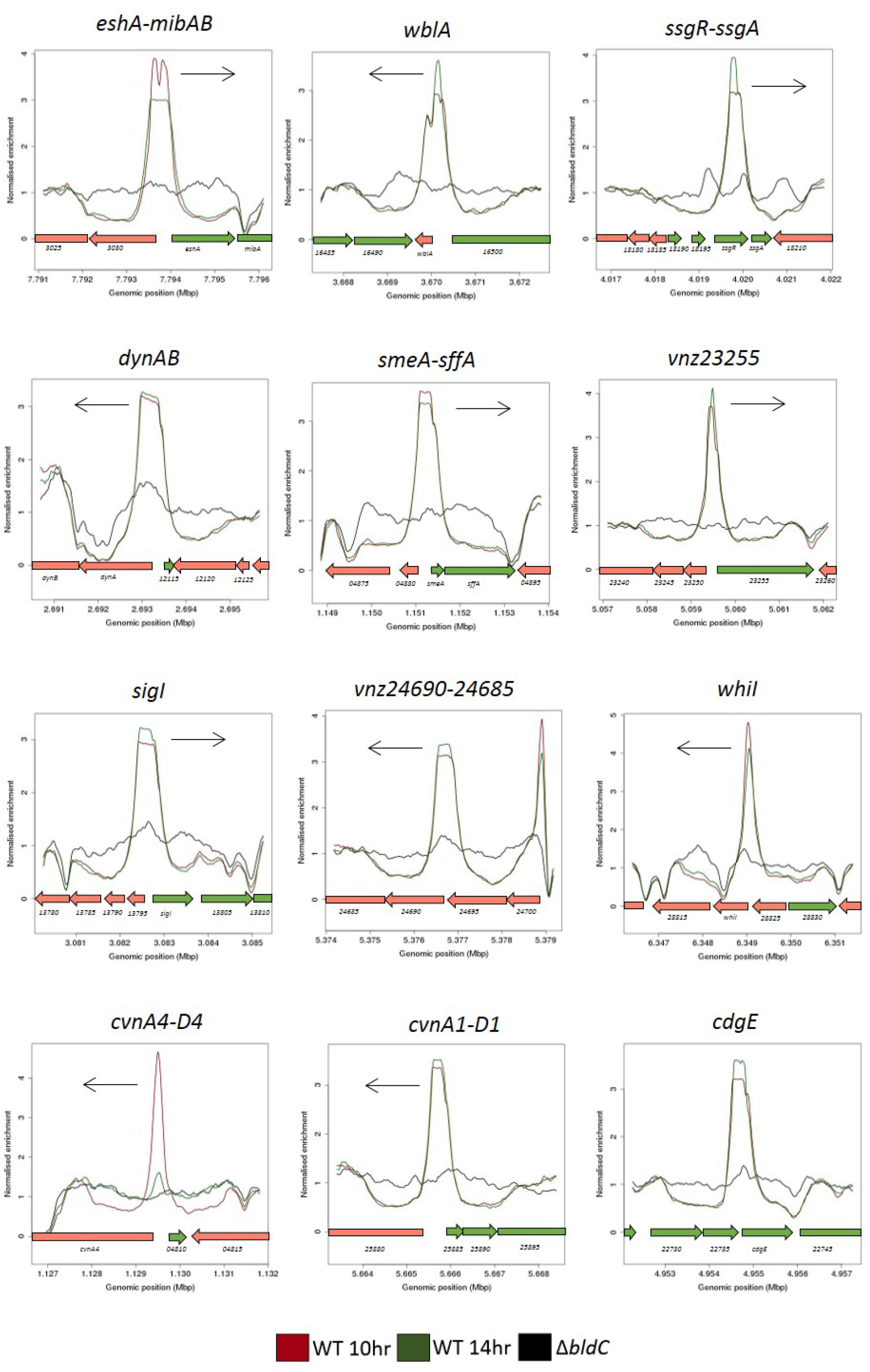
BldC ChIP-seq in *S. venezuelae*. ChIP traces are shown for 12 selected BldC target genes/operons: *eshA-mibAB*, *wblA*, *ssgR-ssgA*, *dynAB*, *smeA-sffA*, *vnz23255*, *sigI, vnz24690-24685*, *whiI*, *cvnA4-D4*, *cvnA1-D1* and *cdgE*. Color-coding of the ChIP samples is as follows: *S. venezuelae* wild type 10 hr (WT 10 hr, red), *S. venezuelae* wild type 10hr (WT 14 hr, green) and Δ*bldC* mutant 14 hr (Δ*bldC*, black). Plots span approximately 5 kb of DNA sequence. Genes running left to right are shown in green, and genes running right to left are shown in red. The black arrow indicates the gene of interest subject to BldC regulation.

**Table 1.** Selected BldC ChIP-seq targets. Listed is the minimum p-value for the ChiP-seq peak, the gene and the gene product. For each target, the RNA-seq data, showing the relative expression values (logFC) for the Δ*bldC* mutant compared to the wild type at the 10 hr and 14 hr time points is also listed. Significant increases in relative expression (>1) are indicated in red. Significant decreases in relative expression (<-1) are indicated in yellow. Where BldC binding is likely to exert control over multiple genes in a single operon, the data for these genes is also listed.

| min apv | Gene | Product | LogFC 10hr | LogFC 14hr |
| --- | --- | --- | --- | --- |
| 2.42E-10 | vnz_18945 | BldC | - | - |
| 3.03E-18 | vnz_35035 | EshA | -8.1457 | -5.3406 |
|  | vnz_35040 | MibA | -4.8905 | -2.8033 |
|  | vnz_35045 | MibB | -6.4229 | -5.1941 |
| 3.19E-17 | vnz_16495 | WblA | -4.3602 | -5.5303 |
| 1.4E-06 | vnz_27205 | WhiH | -4.2208 | -3.7707 |
| 1.33E-07 | vnz_22005 | BldM | -2.6717 | -3.0249 |
| 5.39E-22 | vnz_18205 | SsgA | -1.8093 | -0.5592 |
|  | vnz_18200 | SsgR | -2.2995 | -2.3179 |
| 4.48E-32 | vnz_04805 | CvnA4 | -0.1815 | -2.1394 |
|  | vnz_04800 | CvnB4 | 0.0016 | -2.6155 |
|  | vnz_04795 | CvnC4 | 0.1926 | -2.5418 |
|  | vnz_04790 | CvnD4 | -0.1034 | -2.7909 |
| 3.37E-15 | vnz_25880 | CvnA1 | 0.0094 | -1.1536 |
|  | vnz_25875 | CvnB1 | -0.0808 | -1.5193 |
|  | vnz_25870 | CvnC1 | -0.1607 | -2.2575 |
|  | vnz_25865 | CvnD1 | -0.0979 | -2.2674 |
| 1.33E-07 | vnz_22000 | WhiD | 6.8916 | 2.4359 |
| 1.33E-15 | vnz_18620 | SigF | 4.4232 | 3.8513 |
| 2.52E-07 | vnz_12110 | DynA | 3.0967 | 2.3950 |
|  | vnz_12105 | DynB | 3.1842 | 2.1040 |
| 1.23E-12 | vnz_04885 | SmeA | 2.7343 | 0.1379 |
|  | vnz_04890 | SffA | 2.7046 | 0.3368 |
| 2.33E-12 | vnz_05545 | SsgB | 2.5471 | 0.7373 |
| 4.06E-09 | vnz_13800 | SigI | 2.5392 | 2.7563 |
| 1.48E-09 | vnz_24690 | FtsW-like | 2.4867 | 1.8583 |
|  | vnz_24685 | penicillin-binding protein | 2.1361 | 2.0431 |
| 2.5E-15 | vnz_22740 | CdgE | 2.2747 | 2.0130 |
| 1.87E-27 | vnz_28820 | WhiI | 1.9805 | 3.0348 |
| 1.22E-35 | vnz_12970 | penicillin-binding protein | 1.8746 | 1.6578 |
| 7.74E-05 | vnz_25950 | HupS | 1.8591 | 1.4814 |
| 7.17E-25 | vnz_23255 | penicillin-binding protein | 1.4519 | 1.4785 |
| 1.04E-17 | vnz_15130 | penicillin-binding protein | 0.8151 | 0.7076 |
| 4.46E-06 | vnz_13645 | WhiB | 0.6179 | 0.5096 |

Schumacher et al. (11) showed that BldC binds to DNA in a head-to-tail fashion at a variable number of direct repeats. So, for example, in the *whiI-*BldC structure, there are two direct repeats resulting in the head-to-tail oligomerisation of two BldC monomers, whereas in the *smeA-*BldC structure there are four direct repeats resulting in the head-to-tail oligomerisation of four BldC monomers. In line with this, our data show a broader BldC ChIP-seq peak at the *smeA* promoter compared to the *whiI* promoter (Fig. 5). Indeed, regions of BldC enrichment across the *S. venezuelae* genome were often noticeably broad. Approximately 60% of BldC targets were defined by narrow, *whiI-*like ChIP-seq peaks, but the remaining ~40% showed ChIP-seq peaks at least as broad as the peak observed at the *smeA* promoter (Fig. S1, Table S2). The cooperative binding of BldC to DNA revealed by structural analysis (11) suggested that dimerization on DNA would be the minimum requirement for DNA binding and that extended multimerization would occur at target promoters carrying additional direct repeats. The BldC-DNA structures identified two major elements that define the specificity of BldC binding: a 4-bp AT-rich sequence with a C or G four-five nucleotides downstream. The consensus direct repeat is 5’-AATT(N_3-4_)(C/G)-3’, but the BldC-*smeA* structure showed that conservation of even this degenerate consensus is not critical for BldC binding. In particular, for the AT-rich sequence, it is the narrowing of the minor groove caused by the AT-rich nature of that sequence that is important, rather than direct base reading by BldC (11). Because of this plasticity, it is not possible to predict BldC binding sites bioinformatically. Nevertheless, using this loose consensus as a guide, it seems likely that the BldC targets with narrow ChIP-seq peaks have two appropriately spaced direct repeat sequences, whereas BldC targets with broad ChIP-seq peaks, such as *smeA-sffA*, *cdgE* and *dynAB*, have more (Fig. S1).

### BldC represses the transcription of a subset of target genes

*whiI* and the *smeA-sffA* operon were originally identified as BldC targets in *S. coelicolor* (11), but ChIP-seq analysis showed that they are also BldC targets in *S. venezuelae* (Fig. 5, Table 1). To assess the regulatory influence of BldC on the *whiI* and *smeA* promoters, we performed qRT-PCR using RNA isolated from wild-type *S. venezuelae* and the *bldC* mutant during vegetative growth (10 hr), when BldC is abundant in wild-type cells (Fig. 6). Under these conditions, expression of both *whiI* and *smeA* is significantly higher in the *bldC* mutant compared to the wild type. This suggests that BldC functions to repress the transcription of these developmental target genes during vegetative growth, consistent with the premature initiation of development seen in a *bldC* mutant. To gain a global view of the regulatory impact of BldC, we conducted RNA-seq to compare the transcriptomes of wild-type *S. venezuelae* and the *bldC* mutant at 10 hr and 14 hr timepoints, when both strains were still growing vegetatively (Table S1B). The RNA was prepared from the same cultures used to make protein extracts for the BldC Western blotting shown in Fig. 4. In line with the qRT-PCR data, *whiI* and *smeA* showed significant increases in expression in the *bldC* mutant compared with the wild type. The *smeA* and *sffA* genes form an operon and, consistent with this, both genes showed similar upregulation of expression at the 10 hr and 14 hr time points (Table 1). In total, 156 of the genes we identified as BldC targets in ChIP-seq showed a greater than 2-fold increase in expression in the *bldC* mutant (Table S1C and Fig. 7). These included the key developmental genes *sigF*, *whiD* and *hupS*, which showed a greater than two-fold increase in expression (logFC >1) in the *bldC* mutant, compared to the wild type at both the 10 hr and 14 hr timepoints (Table 1). qRT-PCR confirmed the upregulation of *sigF*, *whiD* and *hupS* expression in the *bldC* mutant relative to the wild type, as was observed for *whiI* and *smeA* (Fig. 6).

**FIG 6.**
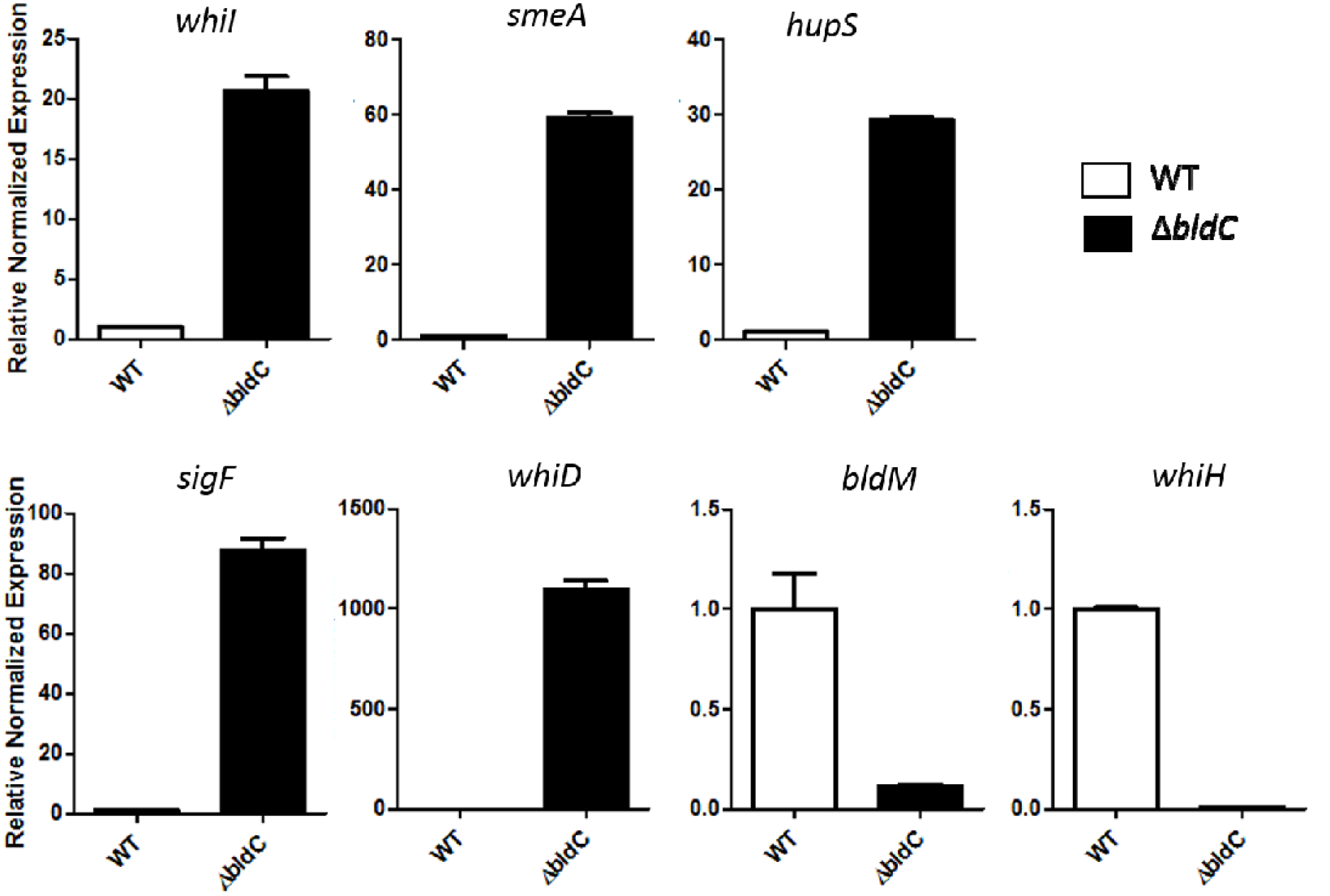
qRT-PCR data showing mRNA abundance for the BldC target genes *whiI, smeA, sigF*, *whiD*, *hupS*, *whiH* and *bldM* in the wild type (white bars) and the *bldC* mutant (black bars). Strains were grown in MYM liquid medium. Expression values were calculated relative to the accumulation of the constitutively expressed *hrdB* reference mRNA and normalized to the wild type value at 10 h.

**FIG 7.**
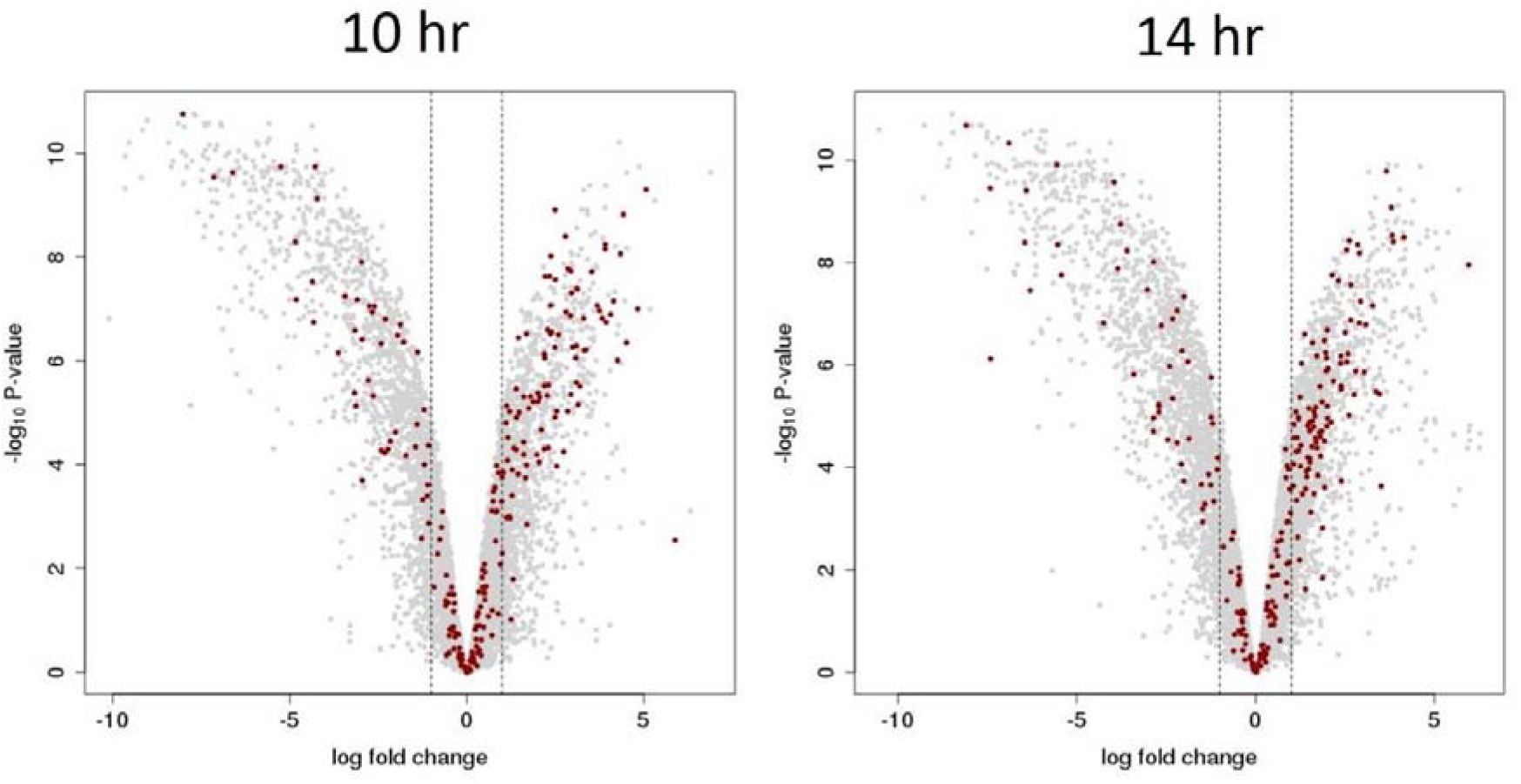
Volcano plots of the RNA-seq data at the 10 hr (left panel) and 14 hr (right panel) time points with significance (-log_10_ P-value) plotted against differential expression (log fold change). The thresholds for significant differential expression (>1 or <-1 log fold change) are indicated via vertical dashed lines. Genes with log fold change >1/<-1 show at least a two-fold increase/decrease in expression in the Δ*bldC* mutant relative to the wild type. Genes that are BldC ChIP-seq targets in *S. venezuelae* are indicated by red dots.

Among the other BldC targets identified by ChIP-seq were a number of genes encoding members of the Penicillin-Binding Protein (PBP) family, required for the synthesis of peptidoglycan (19, 20). Our data indicates that the *vnz12970* and *vnz23255* genes, encoding Class A high molecular mass (HMM) PBPs and the *vnz15130* gene, encoding a low molecular mass (LMM) PBP, are all targets of BldC (Table 1). Expression of these PBP-encoding genes is significantly upregulated in a *bldC* mutant during vegetative growth compared to the wild type, showing that BldC functions to repress their transcription (Table 1).

Peptidoglycan synthesis is required to produce new wall material during cell elongation and to produce septa during division (21). Most rod-shaped bacteria possess distinct gene-pairs to control these two processes, a protein of the SEDS (shape, elongation, division, and sporulation) family and its cognate class B PBP. In *E. coli*, the RodA-PBP2 and FtsW-FtsI pairs control elongation and division respectively (22). In *S. venezuelae*, there are four equivalent SEDS-PBP pairs and our data indicate that one of these pairs, *vnz24690* and *vnz24685*, is under BldC control (Fig. 5 and Table 1). BldC binds upstream of this gene pair and both genes show >4-fold increase (logFC >2) in expression in the *bldC* mutant compared to the wild type.

In *Streptomyces,* the *ftsW-ftsI* SEDS-PBP gene pair is specifically required for cell division at sporulation septa (23). Both genes are found in the division and cell wall (*dcw*) gene cluster. This cluster encodes many proteins that play critical roles in hyphal polar growth, peptidoglycan biosynthesis and cell division, including DivIVA, SepF, SepG, FtsW and FtsZ. Closer inspection of the ChIP-seq data showed that BldC binds at multiple positions across the *dcw* cluster (Fig. 8), although these peaks all fall just below the significance threshold we applied (p < E-04) (Table S1A). The majority of genes within this cluster show a modest increase in expression (logFC 0.5-1.5) during vegetative growth in the *bldC* mutant relative to the wild type (Table S1D), suggesting that BldC functions to repress the transcription of genes within the *dcw* cluster during vegetative growth, in line with the premature initiation of cell division during sporulation that we observe in a *bldC* mutant.

**FIG 8.**
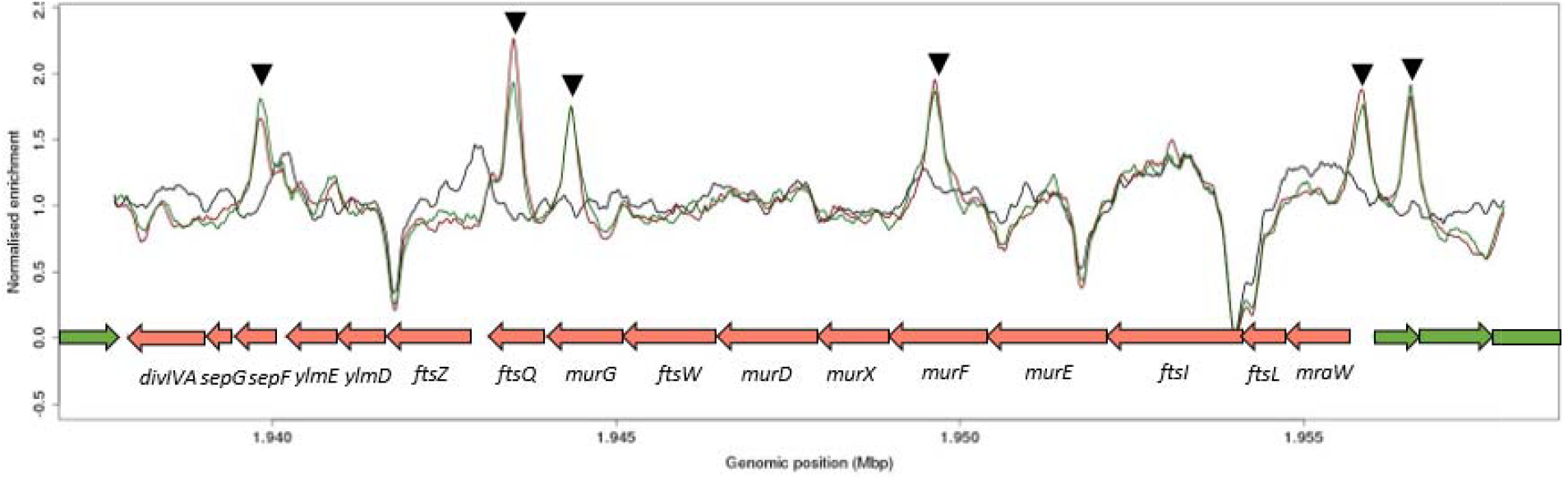
BldC binds at multiple positions across the *dcw* operon. The genes found in the *dcw* cluster are annotated. Color-coding of the ChIP samples is as follows: *S. venezuelae* wild type 10 hr (WT 10 hr, red), *S. venezuelae* wild type 10 hr (WT 14 hr, green) and Δ*bldC* mutant 14 hr (Δ*bldC*, black). Genes running left to right are shown in green, and genes running right to left are shown in red. The black arrows indicate BldC binding sites identified in this analysis.

Our data also indicate BldC-mediated repression of other genes with critical roles in cell division and sporulation such as the *dynAB* operon and *ssgB* (Fig. 5, Table 1). *dynA* and *dynB* encode two dynamin-like membrane-remodelling proteins that stabilize FtsZ rings during sporulation septation via protein-protein interactions with other divisome components including FtsZ, SepF, SepF2 and SsgB (13). In *S. coelicolor,* the actinomycete-specific proteins SsgA and SsgB positively control the spatial distribution of FtsZ rings during sporulation-specific cell division. SsgA binds and recruits SsgB, which in turn recruits FtsZ, determining the future sites of sporulation septation (24). Our data indicate that *ssgB* is a target of BldC-mediated repression (Table 1).

### BldC activates the transcription of a subset of target genes

Strikingly, our RNA-seq data also reveal that large numbers of genes are significantly downregulated in the *bldC* mutant during vegetative growth (Table S1B). Many of these genes are not direct BldC targets but nevertheless encode proteins important for the formation of an aerial mycelium, consistent with the bypassing of aerial hypha formation in the *bldC* mutant (Table S1D). For example, for aerial hyphae to break surface tension and grow into the air, they must be covered in an extremely hydrophobic sheath that is composed of two families of developmentally regulated proteins, the chaplins and the rodlins. In wild-type *S. venezuelae*, expression of the *chp* and *rdl* genes is activated at the onset of development, both on plates and during submerged sporulation (14). In contrast, the *chp* and *rdl* genes were not activated during submerged sporulation in the *bldC* mutant (Table S1E).

In addition to these indirect effects, 91 direct BldC target genes showed a greater than 2-fold reduction in expression (logFC <-1) in the *bldC* mutant compared to the wild type during vegetative growth, implying that BldC functions as an activator of these genes (Table S1E and Fig. 7). Two of these BldC target genes encode the developmental regulators BldM and WhiH, both of which showed significant down-regulation in the absence of *bldC* during vegetative growth in the RNA-seq data (Table 1), a result confirmed by qRT-PCR (Fig. 6). Therefore, BldC functions to activate the transcription of *bldM* and *whiH*, which contrasts with its repression of other key developmental genes (e.g. *whiI, smeA, sigF*, *whiD* and *hupS*) and the observed premature initiation of development in a *bldC* mutant. Our ChIP-seq data coupled with our RNA-seq data also suggest that BldC binds and activates the transcription of *ssgA* and *ssgR*, the latter encoding the sporulation-specific activator of *ssgA* (25) (Fig. 5 and Table 1).

Other noticeable targets of BldC-mediated activation include members of a family of highly conserved operons, known as the “conservons”. Each conservon (*cvn*) consists of four or five genes encoding proteins that, based on biochemical studies of Cvn9, may collectively form complexes of proteins at the membrane with roles in signal transduction (26). Our RNA-seq data indicate that each of the seven conservons present on the *S. venezuelae* chromosome (*cvn1, cvn2, cvn3, cvn4, cvn5, cvn7* and *cvn9*) show a significant reduction in expression during vegetative growth in the *bldC* mutant compared to the wild type (Table S1D). Two (*cvn1* and *cvn4*) are direct targets of BldC, as determined by ChIP-seq (Fig. 5, Table 1). The promoter upstream of *cvn1* is also bound by WhiAB (17) and a *cvn1* mutant of *S. coelicolor* is impaired in aerial mycelium formation (27), collectively suggesting that Cvn1 (and perhaps other members of the conservon family) may play a significant but as yet undefined role in *Streptomyces* differentiation.

The importance of cyclic-di-GMP (c-di-GMP) in the control of *Streptomyces* differentiation became clear with the discovery that engineering high levels of this nucleotide second messenger blocks entry into development, resulting in a classic bald phenotype, whereas engineering low levels of ci-di-GMP causes precocious hypersporulation (5, 9). These phenotypes arise, at least in part, because the ability of the master repressor, BldD, to dimerize and repress a suite of sporulation genes during vegetative growth depends on binding to c-di-GMP (5, 9, 28, 29). c-di-GMP metabolism therefore plays a critical role in coordinating entry into reproductive growth. c-di-GMP is synthesized from two molecules of GTP by diguanylate cyclases (DGCs) and our ChIP-seq and RNA-seq data show that expression of *cdgE*, encoding a predicted DGC, is directly activated by BldC (Fig. 5 and Table 1). *cdgE* is present in at least 90% of *Streptomyces* strains, suggesting the role of this DGC will be widely conserved in the genus (30).

*Streptomyces sp.* are noted producers of the terpene 2-methoisoborneol (2-MIB), one of the volatiles that give soil its characteristic earthy odour. Our RNA-seq data show that expression of the genes required for 2-MIB biosynthesis (*mibA-mibB*) was significantly reduced in the *bldC* mutant compared to the wild type (Table S1B). Unexpectedly, the *mibA*-*mibB* genes were found to form an operon with *eshA*. *eshA* encodes a putative cyclic nucleotide-binding protein of unclear function that is not required for the biosynthesis of 2-MIB (31-33). The effect of BldC on *mibAB* expression is direct. ChIP-seq analysis showed that BldC binds to the promoter of the *eshA-mibA-mibB* operon (Fig. 5 and Table 1) and that all three genes show a similar reduction in expression in the *bldC* mutant, indicating that BldC serves to activate transcription of the operon (Table 1).

## DISCUSSION

Canonical *bld* mutations block entry into development and so the resulting colonies do not form aerial hyphae and spores. These mutations typically define positive regulators such as the response regulator BldM (16) or the sigma factor BldN (14). Although our data indicate that BldC can function as both an activator and a repressor, we have shown that *S. venezuelae bldC* mutants are bald because they enter development prematurely, bypassing the formation of aerial hyphae, and that this correlates with premature expression of a subset of BldC target genes with roles in *Streptomyces* differentiation. Thus, phenotypically, BldC functions as a repressor to sustain vegetative growth and delay entry into development. As such, BldC joins a growing class of Bld regulators known to function as a developmental “brake” (8).

BldD was the first Bld regulator of this alternative class to be clearly recognized. BldD sits at the top of the developmental cascade and represses a large regulon of ~170 sporulation genes during vegetative growth. BldD activity is controlled by the second messenger c-di-GMP, which mediates dimerization of two BldD protomers to generate a functional repressor. In this way, c-di-GMP signals through BldD to repress expression of the BldD regulon, extending vegetative growth and inhibiting entry into development (5, 9, 28, 29). Because a BldD-(c-di-GMP) complex represses the BldD regulon and not BldD alone, engineering the degradation of c-di-GMP *in vivo* also causes a precocious hypersporulation phenotype like that of a *bldD* null mutant (9).

More recently, *bldO* was identified as a second member of this emerging class of *bld* mutant (7, 8). In contrast to BldD and BldC, which both control large regulons, BldO functions to repress a single developmental gene, *whiB*. The precocious hypersporulation phenotype of the *bldO* mutant arises from premature expression of *whiB*, and in line with this, constitutive expression of *whiB* alone is sufficient to induce precocious hypersporulation in wild-type *S. venezuelae* (7). WhiA and WhiB act together to co-control the same set of promoters to initiate developmental cell division in *Streptomyces* (15, 17). WhiA is constitutively present throughout the life cycle, but it only binds to its target promoters at the onset of sporulation when WhiB is present (15, 17). This is because WhiA and WhiB function cooperatively and *in vivo* DNA binding by WhiA depends on WhiB, and *vice versa* (17). As a consequence, the regulation of *whiB* expression is key in controlling the switch between hyphal growth and sporulation. This critical role for WhiB is reflected in the extensive developmental regulation to which *whiB* transcription is subject, being directly repressed by BldC, BldD (28) and BldO (7), and directly activated by BldM (16).

BldC-family members radiate throughout the bacterial domain. Interestingly, some BldC orthologs are annotated as possible DNA resolvase/integrase-associated proteins, consistent with the structural similarity observed between BldC and Xis (11, 34). Xis is a DNA architectural protein that mediates the formation of a nucleoprotein complex required for the phage-encoded Int recombinase/integrase to catalyse the site-specific recombination event that leads to the excision of phage lambda from the *E. coli* chromosome. Like BldC, Xis binds to DNA in a head-tail fashion to generate a nucleoprotein filament, leading to distortion of the DNA (34). Thus, BldC may represent an evolutionary link between transcription factors of the MerR family and DNA architectural proteins (11).

There is an interesting analogy between the relationship of BldC to MerR and the relationship of Fis to NtrC. Fis is a 98-residue nucleoid-associated protein found in proteobacteria that is closely related to the DNA-binding domain of the much larger bacterial enhancer-binding protein NtrC (35-37). Like BldC, Fis prefers binding to A+T-rich DNA and its interaction with DNA is affected by the width of the minor groove (38). Fis can function in the cell as an architectural protein in the nucleoid, but it can also function as a transcription factor (39, 40). Like BldC, Fis exerts a global influence on the transcription profile of the cell and can have positive or negative effects on the activity of its target promoters (41). Fis does not bind a ligand and it is not known to be controlled by post-translational modification. Instead, its influence appears simply to reflect Fis protein concentration, which is high in early log phase but low at other growth stages. In the future, it will be interesting to determine if the activity of BldC is controlled post-translationally, or whether BldC function is more akin to that of nucleoid-associated proteins like Fis.

## MATERIALS AND METHODS

### Construction and complementation of an *S. venezuelae bldC* null mutant

Using ‘Redirect’ PCR targeting (42, 43), *bldC* mutants were generated in which the coding region was replaced with a single apramycin resistance (*apr*) cassette. A cosmid library that covers > 98% of the *S. venezuelae* genome (M.J. Bibb and M.J. Buttner, unpublished) is fully documented at http://strepdb.streptomyces.org.uk/. Cosmid 4O24 was introduced into *E. coli* BW25113 containing pIJ790 and the *bldC* gene (*sven3846*) was replaced with the *apr-oriT* cassette amplified from pIJ773 using the primer pairs bldCdis_F and bldCdis_R. The resulting disrupted cosmids were confirmed by restriction digestion and by PCR analysis using the flanking primers bldCcon_F and bldCcon_R, and introduced into *S. venezuelae* by conjugation (44). Null mutant derivatives, generated by double crossing over, were identified by their apramycin-resistant, kanamycin-sensitive and morphological phenotypes, and their chromosomal structures were confirmed by PCR analysis using the flanking primers bldCcon_F and bldCcon_R. A representative *bldC* null mutant was designated SV25. For complementation, *bldC* was amplified with the primers bldCcomp_F and bldCcomp_R, generating an 846bp fragment carrying the coding sequence and the *bldC* promoter, and cloned into HindIII-KpnI/Asp718 cut pIJ10770 to create pIJ10618. The plasmid was introduced into the *bldC* mutant by conjugation and fully complemented all aspects of the mutant phenotype.

### Time-lapse imaging of *S. venezuelae*

Fluorescent time-lapse imaging was conducted essentially as described previously (7, 12, 13). Before imaging, fresh *S. venezuelae* spores for each of the strains imaged were first prepared by inoculating 30 ml cultures of MYM with 10 μl of the appropriate spore stock or 20 μl of the appropriate mycelial culture. Cells were cultured at 30 °C and 250 rpm until fully differentiated (16-24 h for hypersporulating strains, otherwise 36-40 h). 1 ml of each culture was spun briefly to pellet mycelium, the supernatant spores were diluted 1:50 in fresh MYM, and 50 μl was transferred to the cell loading well of a prepared B04A microfluidic plate (Merck-Millipore). The remaining culture was filter-sterilized to obtain spent MYM that was free of spores and mycelial fragments. The ONIX manifold was then sealed to the B04A plate before transferring to the environmental chamber, pre-incubated at 30 °C. Spores were loaded onto the B04A plate, at 4 psi for 15 seconds using the ONIX microfluidic perfusion system. Fresh MYM medium was set to flow at 2 psi during the first 3 hours during germination, before the 2-psi flow of spent MYM medium for the remainder of the experiment. The system was left to equilibrate for 1 h prior to imaging.

Imaging was conducted using a Zeiss Axio Observer.Z1 widefield microscope equipped with a sCMOS camera (Hamamatsu Orca FLASH 4), a metal-halide lamp (HXP 120V), a hardware autofocus (Definitive Focus), a 96-well stage insert, an environmental chamber, a 100x 1.46 NA Oil DIC objective and the Zeiss 46 HE shift free (excitation500/25 nm, emission 535/30 nm) filter set. DIC images were captured with a 150 ms exposure time, YFP images were captured with a 100 ms exposure time. Images were taken every 30 min. In all experiments, multiple x/y positions were imaged for each strain and in each experiment. Representative images were transferred to the Fiji software package (http://fiji.sc/Fiji), manipulated and converted into the movie files presented here, as described previously (12).

## ChIP-seq, RNA-seq, qRT-PCR, Western blotting and scanning electron microscopy

See Supplemental Materials and Methods (Text S1).

## ACKNOWLEDGEMENTS

We are grateful to Ray Dixon and Charlie Dorman for helpful discussion, and to Genewiz for expert handling of the ChIP and RNA samples.

## SUPPLEMENTAL FIGURE AND TABLE LEGENDS

**Text S1.** Supplemental Materials and Methods.

**FIG S1.** BldC ChIP-seq peaks fall into two classes. BldC binding upstream of some targets generates a broad region of enrichment. For *smeA*, this likely corresponds with the binding of four direct repeats by BldC, observed *in vitro* (Schumacher MA, den Hengst CD, Bush MJ, Le TBK, Tran NT, Chandra G, Zeng W, Travis B, Brennan RG, Buttner MJ. Nat Commun. 2018 9:1139. doi:10.1038/s41467-018-03576-3). Other BldC ChIP-seq targets e.g. *cdgE, dynAB* display similarly broad regions of enrichment and examination of the nucleotide sequence in these regions likewise reveals four similar and appropriately spaced direct repeats. At other BldC ChIP-seq targets, much narrower regions of enrichment are observed. For *whiI*, this likely corresponds with the binding of just two direct repeats by BldC, observed *in vitro* (Schumacher MA, den Hengst CD, Bush MJ, Le TBK, Tran NT, Chandra G, Zeng W, Travis B, Brennan RG, Buttner MJ. Nat Commun. 2018 9:1139. doi:10.1038/s41467-018-03576-3). Other BldC ChIP-seq targets e.g. *vnz23255* and *cvnA4-D4* display similarly narrow regions of enrichment and examination of the nucleotide sequence in these regions likewise reveals a pair of similar and appropriately spaced direct repeats. The ChIP-seq panels are identical to those shown in Fig. 5. The AT-rich sequences of the verified (in the case of *whiI* and *smeA*) and candidate direct repeats to which BldC binds are highlighted below in yellow in the 5′-3′ direction.

**Table S1A.** ChIP-seq data set for *S.venezuelae* BldC.

Only those peaks with significance p < E-04 for at least one of the timepoints are included in the analysis. “Pos” = position in the *S. venezuelae* genome in bases. “lndiff” = the difference between the local normalized (ln) values of the immunoprecipitated wild type samples and Δ*bldC* mutant for each of the time points (lndiff 10hr and lndiff 14hr). “min apv” = minimum adjusted p-value for the 10 hr and 14 hr timepoints. “Closestgene” = nearest annotated gene relative to the position of significance. “lgene” = the identity of a gene where present on the left of the significant position and in the 5’-3’ direction. “lproduct” = the predicted gene product of the lgene. “ldist” = the distance between the significant position and the predicted start codon of the lgene. “igene” = the identity of a gene where the significant position is found within (“in”) a coding region. “iproduct” = the predicted gene product of igene. “idist” = the distance between the significant position and the predicted start codon of the igene. “rgene” = the identity of a gene where present on the right of the significant position and in the 5’-3’ direction. “rproduct” = the predicted gene product of the rgene. “rdist” = the distance between the significant position and the predicted start codon of the rgene. Where present, for each of lgene, igene and rgene, the relative expression values generated by RNA-seq are listed for each time point (lgene/igene/rgene logFC 10hr and lgene/igene/rgene logFC 14hr). Where the logFC >1, cell values are highlighted in red, where the logFC<-1, cell values are highlighted in yellow.

**Table S1B.** Complete RNA-seq data for BldC. Shown is the relative gene expression for the Δ*bldC* mutant compared to the wild type at the 10 hr and 14 hr timepoints. For each gene, the log fold change (logFC) and average p-value (apv) is listed at each timepoint. The canonical gene name (if known) and expected gene product are also listed. Where the logFC >1, cell values are highlighted in red, where the logFC<-1, cell values are highlighted in yellow.

**Table S1C.** BldC represses the transcription of genes during vegetative growth. Listed are BldC ChIP-seq targets that are significantly upregulated (logFC >1) in the absence of *bldC* during vegetative growth at either the 10 hr or 14 hr timepoints, as determined by RNAseq. Ordered by logFC at 10hr and then by logFC at 14hr. For each ChIP-seq target, the position, minimum adjusted p-value (min apv), the gene, expected product and distance to the predicted start codon (dist) is also listed. Where the logFC >1, cell values are highlighted in red, where the logFC<-1, cell values are highlighted in yellow.

**Table S1D.** Selected RNA-seq data for the *dcw* cluster (A), the conservons (B) and genes involved in aerial mycelium formation (C) For each gene, the gene product and log fold change (logFC 10 hr and logFC 14 hr) is listed. Where the logFC >1, cell values are highlighted in red, where the logFC<-1, cell values are highlighted in yellow.

**Table S1E.** BldC activates the transcription of genes during vegetative growth. Listed are BldC ChIP-seq targets that are significantly downregulated (logFC <-1) in the absence of *bldC* during vegetative growth, as determined by RNAseq. Ordered by logFC at 10hr and then by logFC at 14hr. For each ChIP-seq target, the position, minimum adjusted p-value (min apv), the gene, expected product and distance to the predicted start codon (dist) is also listed.

**Table S2.** Analysis of BldC ChIP-seq “peaks”. For each maximum significant position (Max Sig Pos) or ChIP-seq “peak” (as listed in Table S1A), the genomic positions of the left-most (Pos L) and right-most (Pos R) significant positions are recorded as well as the distance between these positions (Width) at both the 10 hr and 14 hr timepoints. “FASTA” = nucleotide sequences between pos L and pos R. “Nearest Gene” = closest gene to the maximum significant position. “Class” = the class of BldC enrichment observed upon manual inspection of the ChIP-seq peak. BldC binding generates either a narrow or broad region of enrichment. BldC targets with broad regions of enrichment generally correlate to “peak” widths >300bp and examination of the nucleotide sequences reveals multiple AT-rich sequences that would support BldC-multimerisation similar to that observed in the *smeA*-BldC structure (Schumacher MA, den Hengst CD, Bush MJ, Le TBK, Tran NT, Chandra G, Zeng W, Travis B, Brennan RG, Buttner MJ. Nat Commun. 2018 9:1139. doi:10.1038/s41467-018-03576-3). BldC targets with narrow regions of enrichment generally correlate to “peak” widths <300bp and examination of the nucleotide sequences reveals fewer AT-rich sequences that would support BldC-multimerisation similar to that observed in the *whiI*-BldC structure (Schumacher MA, den Hengst CD, Bush MJ, Le TBK, Tran NT, Chandra G, Zeng W, Travis B, Brennan RG, Buttner MJ. Nat Commun. 2018 9:1139. doi:10.1038/s41467-018-03576-3). Peaks recorded as “Broad*” are broad and >300bp upon manual inspection but their significance based upon the threshold applied in our analysis means that a much narrower region is considered bioinformatically.

**Table S3.** Strains, Plasmids and Oligonucleotide primers used in this study

**Movie S1.** Time-lapse microscopy of the wild-type strain carrying the FtsZ-YPet fusion. DIC (A) and YFP-channel (B) movies are at 5 frames per second. The time following the first image is indicated at the bottom left. Images were taken every 30 minutes (DIC 150 ms; YFP 100 ms). Movies were assembled in the Fiji software package (http://fiji.sc/Fiji).

**Movie S2.** Time-lapse microscopy of the *bldC* mutant carrying the FtsZ-YPet fusion. DIC (A) and YFP-channel (B) movies are at 5 frames per second. The time following the first image is indicated at the bottom left. Images were taken every 30 minutes (DIC 150 ms; YFP 100 ms). Movies were assembled in the Fiji software package (http://fiji.sc/Fiji).

**Movie S3.** Time-lapse microscopy of the complemented strain carrying the FtsZ-YPet fusion. DIC (A) and YFP-channel (B) movies are at 5 frames per second. The time following the first image is indicated at the bottom left. Images were taken every 30 minutes (DIC 150 ms; YFP 100 ms). Movies were assembled in the Fiji software package (http://fiji.sc/Fiji).

